# Transcriptome-Inspired Spiking Simulations Uncover Human-Specific Prefrontal Dynamics and Provide a Mechanistic Platform for Species-Appropriate Disease Modeling

**DOI:** 10.64898/2026.08.02.741879

**Authors:** Qian-Quan Sun, Yihan Wang, Chunzhao Zhang

**Affiliations:** Graduate Neuroscience Program, University of Wyoming, Laramie, WY 82071, USA; Department of Zoology and Physiology, University of Wyoming, Laramie, WY 82071, USA; Wyoming Sensory Biology Center of Biomedical Research Excellence, University of Wyoming, Laramie, WY 82071, USA

**Keywords:** Cross-species modeling, transcriptomics, human DLPFC, mouse mPFC, Hodgkin–Huxley model, conductance-based spiking networks, high-frequency oscillations, parvalbumin interneurons, gap junctions, excitation–inhibition balance, Brian2

## Abstract

Whole-brain transcriptomic atlases are now widely available, yet computational neural models are almost exclusively parameterized from rodent data and used to infer human brain function, an extrapolation whose cost remains unquantified. To address this, we constructed a biophysically detailed, conductance-based Hodgkin–Huxley spiking microcircuit of a five-population prefrontal network, where every ion-channel, receptor, and gap-junction conductance was scaled by cell-type-specific gene expression. We parameterized the identical circuit using single-nucleus RNA-seq from mouse mPFC and human DLPFC, alongside a literature-derived baseline, and compared their high-frequency-oscillation (HFO) outputs across seven physiological and pathological states. While population firing rates differed only modestly between the two refinements (∼20% for pyramidal and PV cells), the oscillatory dynamics diverged dramatically. The human-refined circuit generated strongly synchronized PV activity and robust ripple- and fast-ripple-band power (e.g., healthy-wake ripple power, in arbitrary units: 322 vs. 24 and 22), whereas the mouse-refined and literature arms remained asynchronous (interneuron synchrony: 0.21 vs. 0.02). This human ≫mouse ≈ original hierarchy was statistically consistent across all seven states (significant arm differences in 75/77 comparisons). Mechanistically, the human transcriptome drove markedly stronger PV–PV electrical coupling (gap-junction scale: 1.78 vs. 0.96) paired with stronger recurrent pyramidal excitation, which collectively synchronized the fast-spiking PV population into a coherent rhythm that perisomatic inhibition then imposed on the local field potential. Critically, these results are model-dependent; the gene-to-conductance mapping is phenomenological, and mRNA expression does not linearly translate to functional conductance. Nonetheless, under this mapping the divergence localizes PV-mediated coupling and excitation–inhibition balance as the parameters most in need of human-specific recalibration. More broadly, this work establishes transcriptome-informed spiking simulation as a powerful strategy for uncovering species-specific computational principles and for building mechanistically grounded, human-relevant models of prefrontal circuit dysfunction, an approach that moves beyond generic rodent defaults to enable targeted, species-appropriate modeling of neurological and psychiatric disorders.

## Introduction

Single-cell and spatial transcriptomics now profile gene expression at cellular resolution across tissues, species, and conditions, and whole-brain atlases span mouse, human, and non-human primates through initiatives such as the Allen Brain Atlas, the Human Cell Atlas, and the BRAIN Initiative Cell Census Network. These data identify which cell types exist, how they differ molecularly, and how those programs change with age, disease, and treatment. Yet this molecular detail has been only loosely connected to the computational models used to study how the same circuits compute and, when they fail, generate pathological activity.

Several recent studies have begun to close this gap. Nandi et al. (2022) generated 9,200 conductance-based single-neuron models from mouse visual cortex and showed that fitted ion-channel conductances emergently predicted differentially expressed channel genes, including the Kv3.1 (Kcnc1) gradient across interneurons. Bernaerts et al. (2025) fit Hodgkin–Huxley models to Patch-seq recordings from mouse motor cortex and linked gene expression to biophysical parameters (R^2^ = 0.17), and Gupta et al. (2025) built cardiac-neuron models directly from single-cell transcriptomes. These efforts share the premise that cell-type gene expression constrains biophysical parameters, but each remains within a single species.

This single-species anchoring is consequential, because models are typically parameterized from mouse and then used to reason about human brain function. Prefrontal cortex illustrates the risk: mouse and human share conserved cell classes, pyramidal cells and PV, SST, VIP, and CCK interneurons, but assemble them in different proportions and with divergent intrinsic properties. Human supragranular pyramidal neurons are larger, lower-resistance, higher-threshold cells with stronger, more uniform HCN-mediated conductances than their mouse counterparts, and human DLPFC is more interneuron-rich (on a PV+SST+VIP basis, the excitatory/inhibitory cell-count ratio is ≈24.7 in mouse vs ≈3.6 in human, recognizing that cell counts do not equal synaptic E/I). Substituting mouse parameters into a model meant to represent human cortex may therefore misstate excitability and its pathological signatures.

Here we ask how much that substitution costs. Using a transcriptomically informed pipeline that maps cell-type gene expression onto the ion-channel, synaptic, and gap-junction conductances of a five-population, conductance-based Hodgkin–Huxley (HH) prefrontal microcircuit, we parameterize the same circuit from two single-nucleus RNA-seq datasets, mouse mPFC and human DLPFC, and run both through an identical HFO pipeline alongside a literature baseline (Figure 1). We treat this as a methods-and-comparison study: the mapping is phenomenological, so we validate it explicitly, interpret the cross-species contrast qualitatively, and use it to localize the parameters most in need of human recalibration.

**Figure 1.**
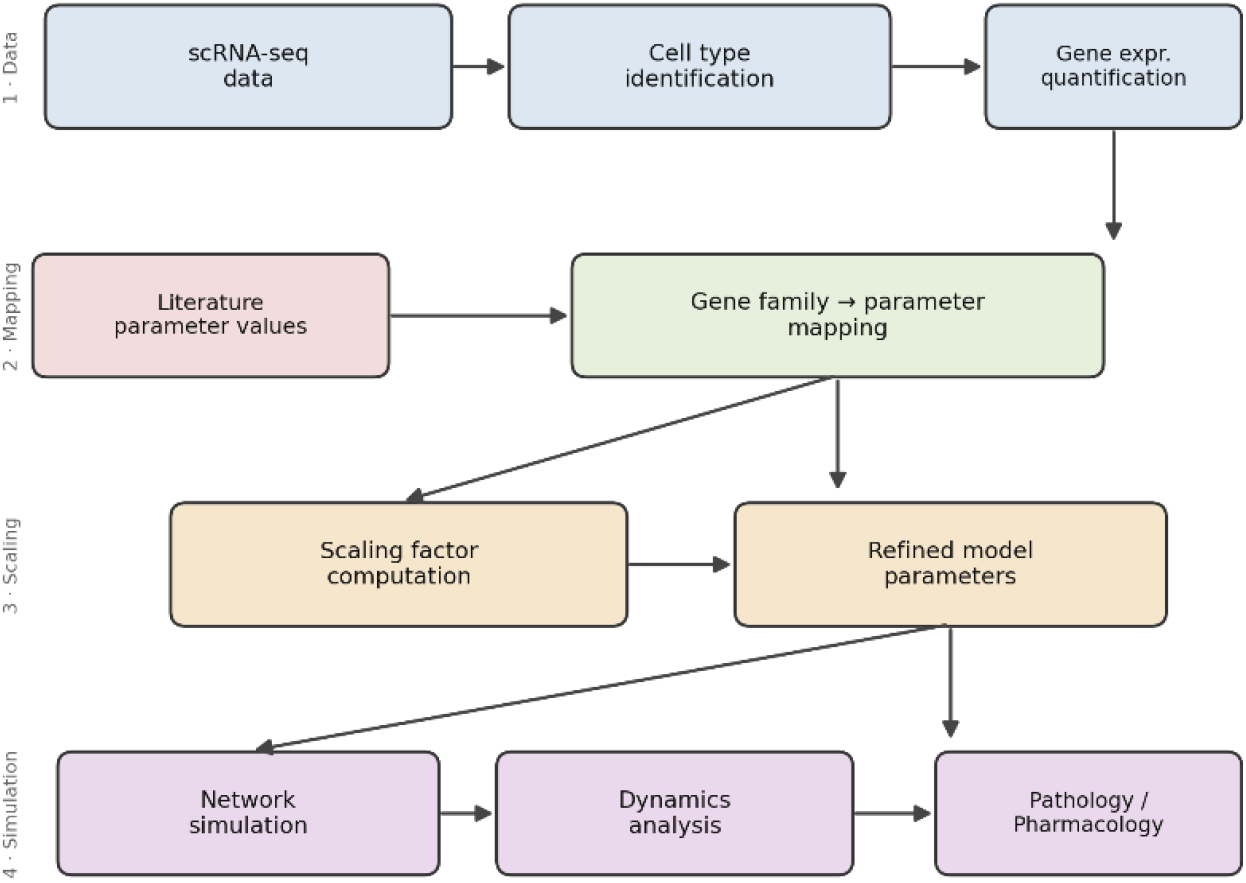
Overview of the transcriptome-to-network pipeline. Single-nucleus transcriptomic data are translated into conductance-based spiking-network parameters, enabling region- and species-specific modeling, disease simulation, and in silico pharmacology.

**Figure 2.**
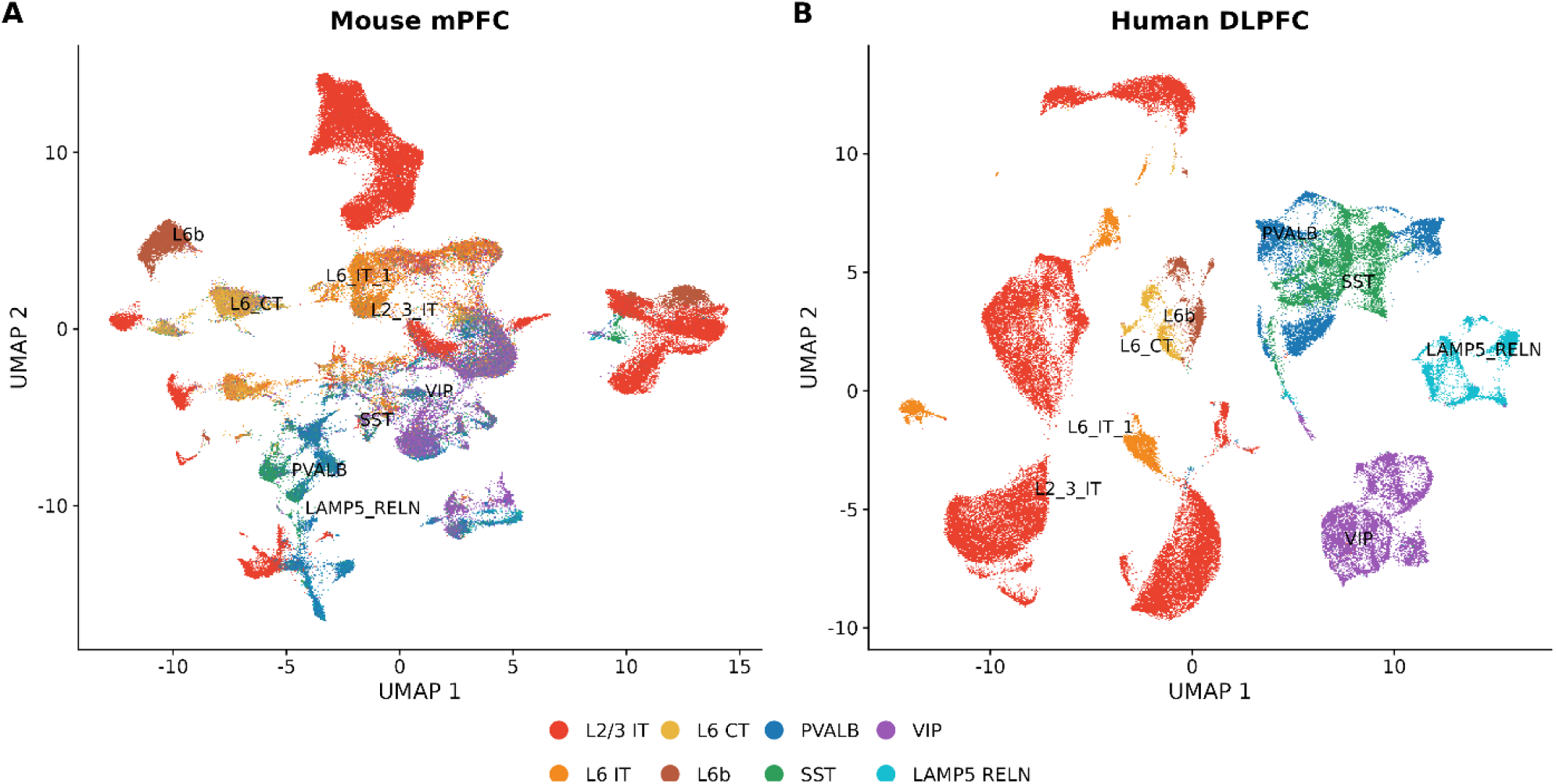
Cross-species UMAP of major cortical neuron subtypes. UMAP embeddings of single-nucleus transcriptomes from mouse mPFC (38,234 neurons) and human DLPFC (GSE213982; 31,727 neurons), processed as described in Sec. 2.1. Points are individual nuclei coloured by annotated cell type. Embeddings were computed separately per species, so the cross-species comparison is qualitative. Both species resolve the conserved excitatory and interneuron classes used to parameterize the five model populations (Pyr, PV, SST, VIP, CCK).

## 1. Methods

The pipeline applies the same steps to each species (Figure 1): single-nucleus RNA-seq is processed to identify cell types and quantify expression; biophysically relevant gene families (ion channels, receptors, connexins) are defined; mean family expression is computed per model cell type; a shared literature baseline conductance is scaled by relative within-species expression; and the resulting conductance-based network is simulated to study high-frequency oscillations. Three parameter arms share identical wiring: a species-neutral literature baseline (original), a mouse-mPFC-refined arm, and a human-DLPFC-refined arm.

### 1.1 Transcriptomic datasets

We parameterized the circuit from two control datasets. The mouse mPFC dataset (Sun Lab scRNA-seq; 3 female controls) yielded 38,234 neurons after quality control, annotated to Allen cell types; the human DLPFC dataset (GSE213982, Maitra et al., 2023; 18 female controls) yielded 31,727 neurons, annotated as layer-resolved excitatory neurons plus PV/SST/VIP/LAMP5/ADARB2 interneurons. Genes were matched across species by uppercasing mouse symbols. Because absolute cross-species levels are confounded by platform and post-mortem interval, all comparisons were made within species after z-scoring, and the cross-species contrast is interpreted qualitatively. The mouse data were generated with micro dissected, acutely isolated mPFC neurons under control conditions, which may bias toward more dendritic expressed transcripts; the human data are from female donors only, a sex confound limiting generalization to male cortex.

### 1.2 Microcircuit architecture

The network comprises 500 single-compartment neurons in five populations, 350 pyramidal (Pyr), 50 parvalbumin (PV), 40 somatostatin (SST), 30 vasoactive-intestinal-peptide (VIP), and 30 cholecystokinin (CCK) cells, connected by conductance-based chemical synapses and within-class electrical synapses. Chemical synapses are conductance-based: each presynaptic class increments a target-specific conductance on the postsynaptic neuron (AMPA for excitation; a separate GABA-A channel per interneuron class), each decaying with its own time constant (AMPA 2 ms; PV 3 ms; CCK 5 ms; SST and VIP 11 ms). Within-class gap junctions couple PV–PV, SST–SST, and VIP– VIP pairs as an ohmic summed current I_gap = Σ w_gap·(V_pre − V_post). Every neuron additionally receives an independent background Poisson excitatory drive that maintains baseline firing; short-term synaptic depression/facilitation and activity-dependent chloride accumulation (a dynamic GABA reversal potential) are included on the relevant pathways. Connection probabilities and target channels are summarized in Table 1.

**Table 1.** **Synaptic connectivity of the microcircuit: connection probability and target channel for each pathway. Synaptic conductances decay with class-specific time constants (AMPA 2 ms; PV 3 ms; CCK 5 ms; SST and VIP 11 ms).**

| Pathway | Prob. | Channel / effect |
| --- | --- | --- |
| Pyr → PV / SST / CCK / VIP | 0.25 | AMPA (feed-forward excitation) |
| PV → Pyr | 0.75 | fast GABA-A (perisomatic) |
| PV → PV | 0.65 | fast GABA-A (recurrent) |
| SST → Pyr | 0.55 | slow GABA-A (dendritic) |
| CCK → Pyr | 0.45 | GABA-A (perisomatic) |
| VIP → SST | 0.65 | GABA-A (disinhibition) |
| VIP → Pyr | 0.10 | slow GABA-A |

### 1.3 Hodgkin–Huxley single-neuron model

Each neuron is a single-compartment membrane governed by eight active and passive currents:

C·dV/dt = I_Na + I_Kdr + I_Kv3 + I_KA + I_Ca + I_KCa + I_M + I_leak + I_syn + I_gap + I_bg (1)

I_Na = gNa·m^3^h·(E_Na − V) and I_Kdr = gKdr·n^4^·(E_K − V): spike generation, Traub–Miles kinetics with a threshold parameter V_T.

I_Kv3 = gKv3·p^4^·(E_K − V): fast, non-inactivating K^+^ that supports high-frequency firing (PV only).

I_KA = gKA·a^4^b·(E_K − V): transient A-type K^+^ (SST/VIP/CCK). I_Ca = gCa·s^2^·(E_Ca − V): high-threshold calcium current.

I_KCa = gKCa·[Ca/(Ca+K_d)]·(E_K − V): calcium-activated K^+^ producing spike-frequency adaptation.

I_M = gM·q·(E_K − V): slow, non-inactivating muscarinic K^+^. I_leak = gL·(E_leak − V): passive leak.

Intracellular calcium is a dynamic variable that decays with time constant τ_Ca and receives a spike-triggered increment Ca_inc; the resulting I_KCa implements the adaptation that distinguishes adapting (SST, CCK, Pyr) from non-adapting (PV) cells. Gating variables follow standard voltage-dependent kinetics; equations are integrated with the stochastic Heun method at dt = 0.05 ms (the background drive contributes an Ornstein–Uhlenbeck noise current). A spike is registered when V crosses −20 mV, followed by a short refractory window, and the action potential repolarizes intrinsically. Per-cell-type baseline conductances were set to reproduce published cortical firing-rate/current (F–I) relationships (Results, Sec. 3.1).

### 1.4 Transcriptome-to-conductance mapping

Cell-type expression tables (rows: the seven sequenced prefrontal types; columns: ion-channel, receptor, and connexin genes) were reduced to gene families and mapped to model conductances (Table 2).

**Table 2.** Mapping of transcriptomic gene families to Hodgkin–Huxley conductances and synaptic parameters.

| Gene family | Genes | Model target |
| --- | --- | --- |
| Na | Scn1a, Scn2a, Scn3a, Scn8a | gNa |
| Kdr | Kcnb1 | gKdr (delayed rectifier) |
| Kv3 | Kcnc1 | gKv3 (fast K<sup>+</sup>; PV) |
| KA | Kcnd2, Kcnd3 | gKA (A-type; SST/VIP/CCK) |
| Ca | Cacna1a, Cacna1b | Ca<sub>inc</sub> (adaptation) |
| Leak | Kcnk1, Kcnk2 | gL |
| AMPA | Gria1, Gria2 | AMPA synaptic weights |
| GABA-A | Gabra1, Gabra2, Gabrb1 | GABA-A synaptic weights |
| Connexin | Gjd2, Gja1, Gjb2 | PV / SST / VIP gap junctions |

For a given cell type and gene family a multiplicative scale factor is computed in five steps: (1) average the family’s genes for that cell type; (2) divide by the average of the same family across a within-species reference set (interneurons for PV/SST/VIP; the four pyramidal laminar subtypes for Pyr); (3) clamp the ratio to [0.5, 2.0]; (4) for Pyr, blend the laminar-subtype factors by prefrontal composition; and (5) apply the factor to the shared literature baseline conductance. Because normalization is within-species, a factor reports how channel-rich a cell type is relative to its own species’ average, the quantity that is comparable across sequencing platforms. gKCa and gM are held at baseline (not gene-scaled). As a worked example, PV’s Na-family expression relative to the interneuron average gives a scale of 1.00 in mouse versus 1.14 in human, human PV cells over-express sodium channels (chiefly Scn1a, encoding Na_v_ 1.1) relative to other human interneurons, whereas mouse PV sits at the interneuron average. The three arms are original (all factors = 1.0), mouse-refined, and human-refined; the largest cross-species differences are in interneuron gap junctions (PV coupling 0.96 mouse vs 1.78 human) and recurrent pyramidal excitation (w_Pyr→Pyr 0.82 vs 1.15). The mapping is phenomenological: pervasive mRNA–protein discordance (reported for 97.5% of genes in at least one neural region; Titlow et al., 2023) and state-dependent mRNA– current uncoupling (Viteri et al., 2024) mean it should be read as a heuristic for relative parameter direction rather than a source of absolute conductances.

### 1.5 Network states and simulation protocol

Three parameter arms, original (literature), mouse-refined, and human-refined, were each simulated over seven conditions × 10 seeds (10 s each, 1 s burn-in), a full sweep of 3 × 7 × 10 = 210 simulations executed in parallel with the numpy code-generation backend. Each run integrates the network at dt = 0.05 ms with a fixed seed and records population firing rates; within-class spike-train synchrony (mean pairwise correlation in 5 ms bins) for pyramidal cells and, pooled, for interneurons; a synthetic LFP sampled at 2 kHz; and ripple/fast-ripple event rates and band powers. The seven states are defined in Table 3.

**Table 3.** Definitions of the seven simulated network states.

| Condition | Definition |
| --- | --- |
| healthy_wake | wake state, all interneurons intact |
| pv_loss | PV output reduced (PV block) |
| vip_high | elevated VIP→SST gain (VIP over-activation / disinhibition) |
| nrem_intact | NREM state, all interneurons intact |
| nrem_pv_low | NREM state with reduced PV output |
| epileptic_nrem | NREM with combined pathology (depolarized GABA reversal, PV↓, VIP↑) |
| gap_on_path | enhanced pyramidal electrical coupling (pyramidal gap junctions on) |

### 1.6 LFP, HFO analysis, and statistics

The LFP is a weighted sum of population spike trains convolved with class-specific kernels (a sharp ∼1 ms pyramidal action-potential kernel and slower inhibitory postsynaptic kernels), giving a 2 kHz signal that resolves the fast-ripple band; it is a simplified surrogate that omits volume conduction, laminar geometry, and dendritic return currents. HFOs are detected by zero-phase Butterworth band-pass filtering (ripple 80–250 Hz, fast ripple 250–500 Hz), taking the Hilbert amplitude envelope, and marking events where the envelope exceeds three times a fixed baseline computed once from a healthy-wake reference, so event rates are comparable across arms and conditions; band power is the integral of the Welch power spectral density over each band. For every condition × metric the three arms were compared by one-way ANOVA when all groups passed a Shapiro–Wilk normality test and by Kruskal–Wallis otherwise, with Tukey HSD or Holm-corrected Dunn post-hoc tests, and bootstrap 95% confidence intervals (1000 resamples, fixed seed) were computed for all pairwise log2 fold-changes between arms. We used α = 0.05 for all hypothesis tests; bootstrap confidence intervals for log_2_ fold-changes used 1,000 resamples across the 10 seeds per condition.

## 2. Results

### 2.1 Single-cell validation

With gene scalings set to 1.0 (literature arm), the five neuron models reproduce the expected cortical firing-rate hierarchy and fall within published F–I ranges. At a 200 pA step, PV cells fire fastest (∼70 Hz, rising steeply with current toward the 80–120 Hz fast-spiking band), while SST, CCK, VIP, and pyramidal cells fire in the ∼8–20 Hz range with clear spike-frequency adaptation in the adapting types. The PV ≫other-type ordering that underlies fast-ripple generation is thus preserved at the single-cell level.

### 2.2 Baseline network dynamics

In the healthy-wake state the network settles into stable, asynchronous-irregular activity with physiological rates (literature arm: pyramidal 23.8 Hz, PV 70.5 Hz, SST 19.3 Hz, CCK 5.0 Hz, VIP 5.0 Hz; Figure 3). Fast-spiking PV cells form the most active population, as required for interneuron-driven high-frequency rhythms.

**Figure 3.**
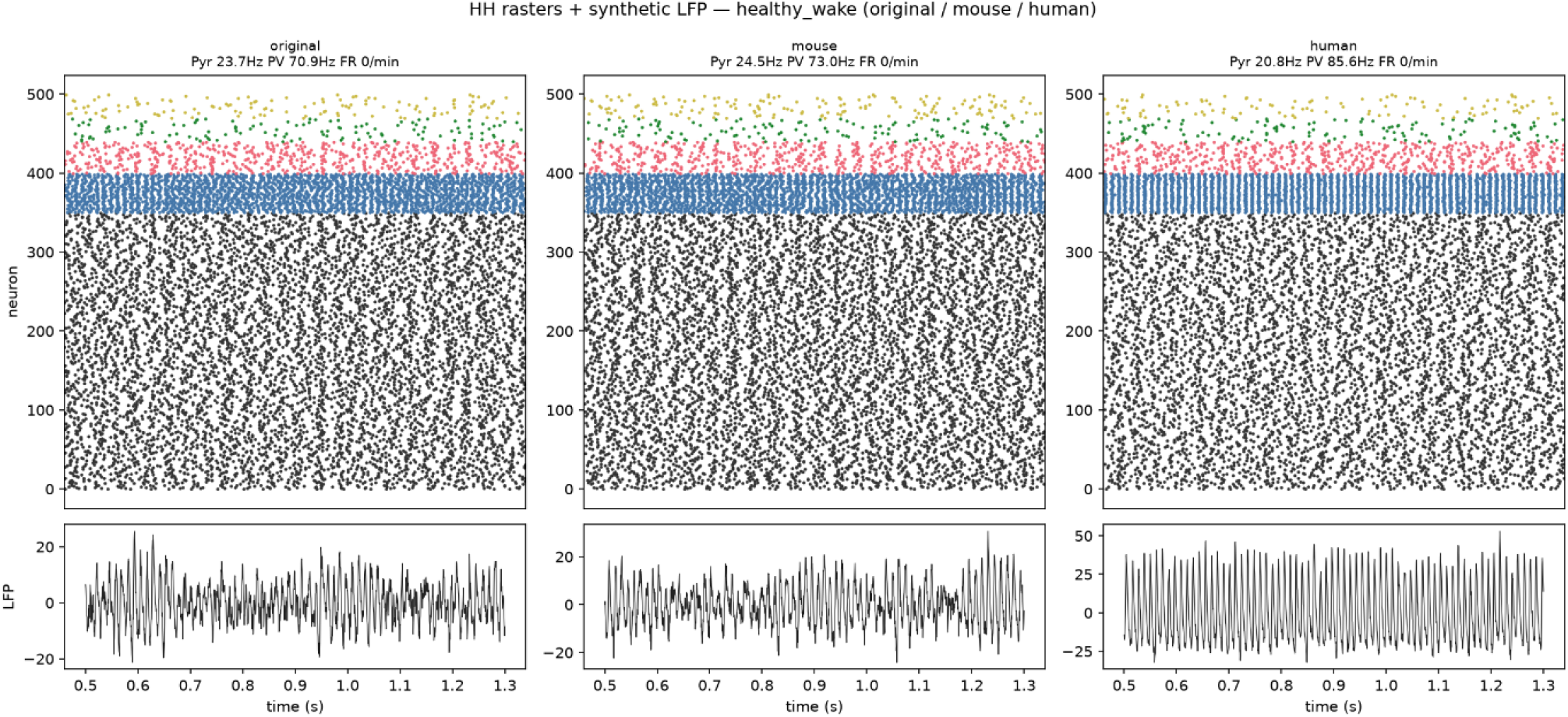
Spike rasters (all five populations, colour-coded) and the synthetic LFP (lower panels) for the three arms in the healthy-wake state. In the human-refined arm the PV band synchronizes and the LFP develops a regular high-frequency oscillation that is absent in the other arms.

### 2.3 Cross-species divergence in high-frequency oscillations

Population firing rates differ only modestly between the mouse- and human-refined arms (pyramidal and PV rates within ∼20% of each other). The striking difference is in the oscillatory regime. In the human-refined arm the interneuron network becomes strongly synchronized and the LFP develops large ripple- and fast-ripple-band power: in healthy wake, ripple-band power is 322 (arb. units) for the human arm versus 24 (mouse) and 22 (original), and interneuron synchrony is 0.21 versus 0.02. All ripple, fast-ripple, and synchrony values reported here are model outputs (band powers in arbitrary units), not experimental measurements, and are interpretable only as comparisons across arms and conditions. This ordering (human ≫mouse ≈ original) holds across all seven network states (Table 4).

**Table 4.** Ripple-band power (arb. units; model output, not an experimental measurement) by network state for the three parameter arms.

| Condition | original | mouse-refined | human-refined |
| --- | --- | --- | --- |
| healthy wake | 22.5 | 24.5 | 322.0 |
| pv loss | 96.8 | 99.8 | 372.3 |
| vip high | 24.6 | 25.5 | 322.6 |
| nrem intact | 10.5 | 9.9 | 62.6 |
| nrem pv low | 15.8 | 15.3 | 87.1 |
| epileptic nrem | 19.1 | 18.3 | 130.6 |
| gap on path | 54.8 | 55.1 | 143.7 |

Fast-ripple-band power and interneuron synchrony follow the same ordering (human ≫mouse ≈ original). The log2 fold-change map (Figure 4) summarizes the contrast across all metrics and conditions: firing rates cluster near zero (little cross-species difference), while every synchrony and HFO-power metric is strongly shifted toward the human arm. The differences are statistically robust, across the 77 condition × metric combinations, 75 showed a significant omnibus difference among arms (p < 0.05), and the mouse-refined versus human-refined post-hoc contrast was significant in 75 of 77 comparisons, and the full sweep of 210 simulations completed with no failed runs.

**Figure 4.**
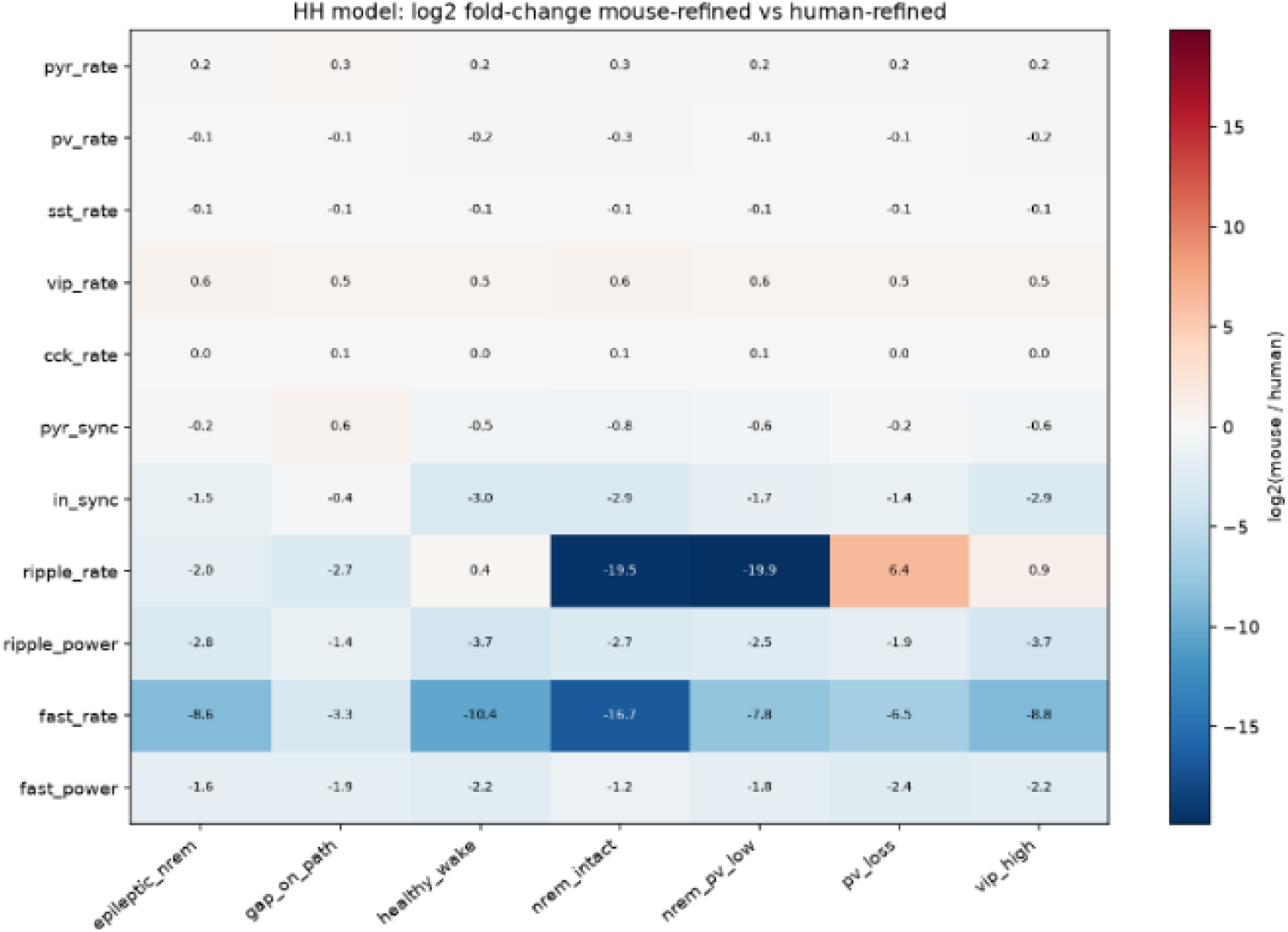
log2 fold-change (mouse-refined / human-refined) of each summary metric across conditions. g rates differ little; ripple/fast-ripple power, event rate, and interneuron synchrony are consistently human-dominated.

**Figure 5.**
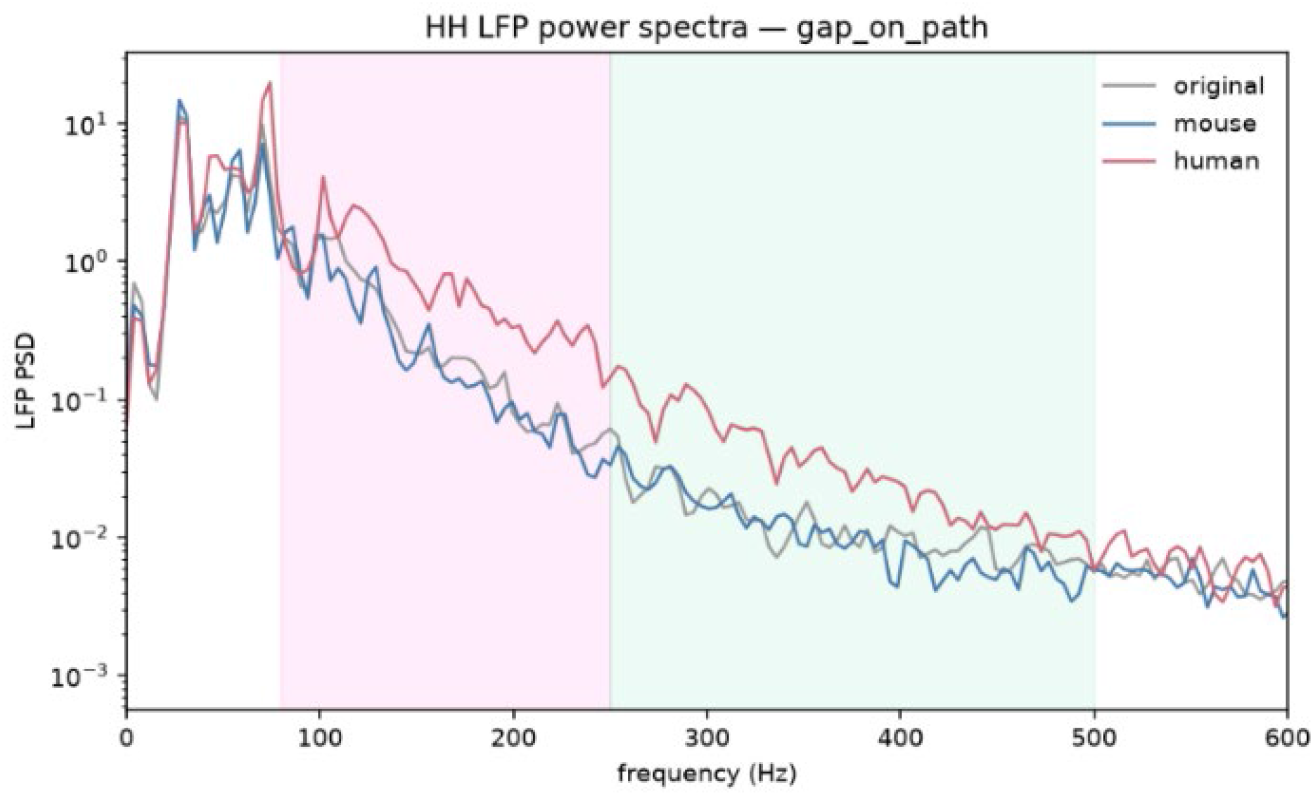
Ripple-band power (model, arbitrary units) by arm (healthy wake); box = interquartile range, points = individual seeds.

### 2.4 Mechanism: PV–PV electrical coupling

The divergence is a network-coupling effect. Under this mapping, the human transcriptome is associated with markedly stronger PV–PV electrical coupling (gap-junction scale 1.78 vs 0.96 in mouse) together with somewhat stronger recurrent pyramidal excitation. Because the PV population fires fast, this enhanced electrical coupling in the model synchronizes the PV population into a coherent fast rhythm, which perisomatic PV→pyramidal inhibition imposes on the pyramidal population and hence on the LFP, producing the elevated ripple and fast-ripple power seen in the human-refined arm. In short, the model predicts that the human-DLPFC interneuron configuration is intrinsically more prone to synchrony-driven high-frequency oscillations than the mouse-mPFC configuration, with the effect concentrated in the PV / fast-ripple regime rather than in mean firing rate.

## 3. Discussion

A human-DLPFC-parameterized Hodgkin–Huxley circuit is markedly more prone to synchrony-driven high-frequency oscillations than the mouse-mPFC-parameterized circuit, even though the two arms differ little in mean firing rate. The effect is driven by stronger PV–PV electrical coupling and somewhat stronger recurrent excitation under the human mapping, which synchronize the fast-spiking PV population into a coherent rhythm that is imprinted on the LFP, whereas the mouse-refined and literature arms remain asynchronous. Transplanting mouse prefrontal parameters into a model meant to reason about human cortex would therefore tend to understate the human circuit’s propensity for PV-synchronized ripple and fast-ripple activity, though the magnitude depends on the phenomenological gene-to-conductance mapping. The divergence also localizes where human recalibration matters most: PV-mediated electrical coupling and excitation–inhibition balance. This is a model-dependent prioritization of which parameters to refine first when human data are scarce, not a claim that all parameters are equally uncertain.

### 3.1 Why the species diverge, and the role of PV electrical coupling

The divergence aligns with the experimental literature: human DLPFC is more interneuron-rich, and human supragranular pyramidal neurons are larger, higher-threshold cells with stronger, more uniform HCN-mediated conductances than mouse. Our within-species mapping cannot resolve absolute conductances, but the direction of the inferred differences, stronger human interneuron coupling and recurrent excitation, concentrates the cross-species effect in the PV network rather than in mean excitability. Ripples and fast ripples are thought to arise from synchronized PV firing and its perisomatic inhibition of pyramidal cells, with interneuronal electrical coupling sustaining that synchrony; our result is a transcriptome-derived instance of this mechanism, in which the inferred human connexin profile supplies the coupling that tips the PV network into synchrony. Because cell-count E/I ratios overstate functional synaptic balance (Loomba et al., 2022), we interpret the composition-linked differences qualitatively.

### 3.2 Evaluation criteria for transcriptome-guided models

As a methods-and-comparison paper, we make explicit four axes on which such models should be judged: single-neuron plausibility (refined firing hierarchies should agree with known relationships), network plausibility (population firing and HFO activity should fall in ranges comparable to existing models), mechanistic consistency (an established mechanism, here PV electrical coupling driving interneuron synchrony, should be expressed coherently rather than manufactured or destroyed across parameterizations), and cross-species specificity (the two parameterizations should diverge in concrete, interpretable ways rather than being interchangeable). The present comparison meets all four: the F–I hierarchy is preserved, network firing rates are physiological, the PV-coupling mechanism is consistent across the seven states, and the two arms diverge in a specific, interpretable regime, PV synchrony and HFO power, that a single-species study cannot test.

### 3.3 Limitations

The gene-to-conductance mapping is phenomenological: mRNA–protein discordance (Titlow et al., 2023) and state-dependent mRNA–current uncoupling (Viteri et al., 2024) mean the scaled conductances recover relative direction, not absolute magnitude, and should not be read as biophysical predictions. Orthologs were matched by symbol uppercasing rather than a curated table; the synthetic LFP omits volume conduction and laminar geometry; and each species rests on a single, female-only dataset, with the micro-dissected single-cell mouse protocol over-representing dendritic mRNAs, so comparisons are within-species, qualitative, and not generalizable to male cortex. Finally, the neurons are single-compartment and AMPA-only, so the PV-coupling mechanism may partly compensate for absent NMDA-mediated slow excitation; isolating the primary driver will require independently manipulating gap coupling, PV inhibition, and NMDA conductance.

### 3.4 Generalizability and future directions

The pipeline is general across species, regions, and tissues; the mouse-versus-human prefrontal comparison is one instance. Priorities follow from the results: recalibrate human PV electrical coupling and E/I balance with targeted human snRNA-seq and, where possible, ex vivo physiology; resolve human IT versus PT pyramidal subtypes with a curated ortholog table; incorporate protein- or electrophysiology-derived constraints to improve the mapping; and add NMDA conductances, male donors, and snRNAseq mouse data to control for sex and capture bias while testing whether PV coupling remains the dominant HFO driver.

### 3.5 Conclusion

The same prefrontal microcircuit can be parameterized from either species’ transcriptome on a common conductance-based substrate, and under our phenomenological mapping the two imply different operating points: the human-DLPFC configuration confers stronger PV–PV electrical coupling and synchronizes into large ripple and fast-ripple oscillations, whereas the mouse-mPFC configuration stays asynchronous and near the literature baseline, with mean firing rates barely distinguishing the arms. Species-specific parameterization is therefore warranted when transcriptome-guided models are used to reason about human prefrontal physiology, and the cross-species comparison turns qualitative species differences into a testable, mechanistic prediction, that human PV electrical coupling is the parameter most in need of recalibration and a primary lever on prefrontal HFO generation. These predictions are model-dependent, so the mapping’s value lies in the cross-species contrast it generates rather than in the absolute values it produces.

### Use of Large Language Models

Large language models (LLMs) were used to help perform the cross-species RNA-seq data analysis, to help with the Python coding, and to help with formatting and proof-reading of the manuscript. LLMs were not used to design the study, generate scientific claims, or interpret the results; all authors reviewed the work and take full responsibility for its content.

## Acknowledgments

We thank the Sun Lab members for helpful discussions. Human data were obtained from GEO accession GSE213982 (Maitra et al., 2023). This work was supported by the National Institutes of Health (R21MH131363, R21MH141703, 2P20GM121310).

## Reproducibility

scRNA-seq comparison in R/Seurat 5; conductance-based Hodgkin–Huxley simulation in Python 3.11 / Brian2 2.9 (human GSE213982; mouse mPFC data from the author’s lab), 3 arms × 7 conditions × 10 seeds = 210 runs. Statistical, robustness, and validation outputs are archived with the model repository.

